# Kelch13 stochastics determine drug survival in resistant malaria parasites

**DOI:** 10.64898/2026.07.13.738300

**Authors:** Isabelle G Henshall, Patricia López-Barona, Sabine Schmidt, Lea Sophie Panitzsch, Vendula Horáčková, Tobias Spielmann

## Abstract

As resistance to the frontline antimalarial artemisinin (ART) in the deadliest malaria species *Plasmodium falciparum* spreads, it threatens the gains made in reducing global malaria burden over the last decades. Mutations in the gene encoding Kelch13 (K13) hold a central role in ART resistance. Yet even in clonal populations harbouring a resistance conferring *k13* mutation, only a portion of parasites survive drug exposure. Here we show that stochastic cell-to-cell variation of cellular K13 levels determines survival of individual parasites. Using isogenic parasite lines, we establish that decreased cellular K13 levels correlate with resistance and in addition a fitness cost through an increased cell cycle length. As resistance increases, parasites show a shift of the stochastic range towards lower K13 levels, increasing the proportion of drug survivors. While this range was inherited, individual parasites with differing K13 levels gave rise to progeny across the full spectrum. Hence, while the resistance fitness profile of each parasite line is defined by the range of K13 levels, the fate of individual parasites within that range is determined by stochastics. These findings provide an explanation why only a portion of genetically identical parasites survive drug exposure and reveal a system where the stochastics of a single protein determine individual parasite drug survival and fitness.

## Introduction

Malaria results in over half a million deaths each year with concern this number will rise with the spread of resistance to the front-line antimalarial, artemisinin and its derivates (ART)^1,6,7^. Mutations in Kelch13 (K13) in *Plasmodium falciparum,* the species responsible for majority of malaria mortality, reduce K13 cellular abundance and reduced haemoglobin uptake rates^2,4,5,8,9^. The resulting reduction in subsequent haemoglobin digestion by-product availability is thought to reduce activation of ART, leading to resistance (ART-R)^10^. Reduction in haemoglobin uptake rates is only an effective resistance strategy in ring stage parasites, the first stage of the 48 hour blood stage lifecycle (Fig. 1a), as subsequent stages have accumulated extensive haemoglobin digestion by-products that activate ART.

**Figure 1.**
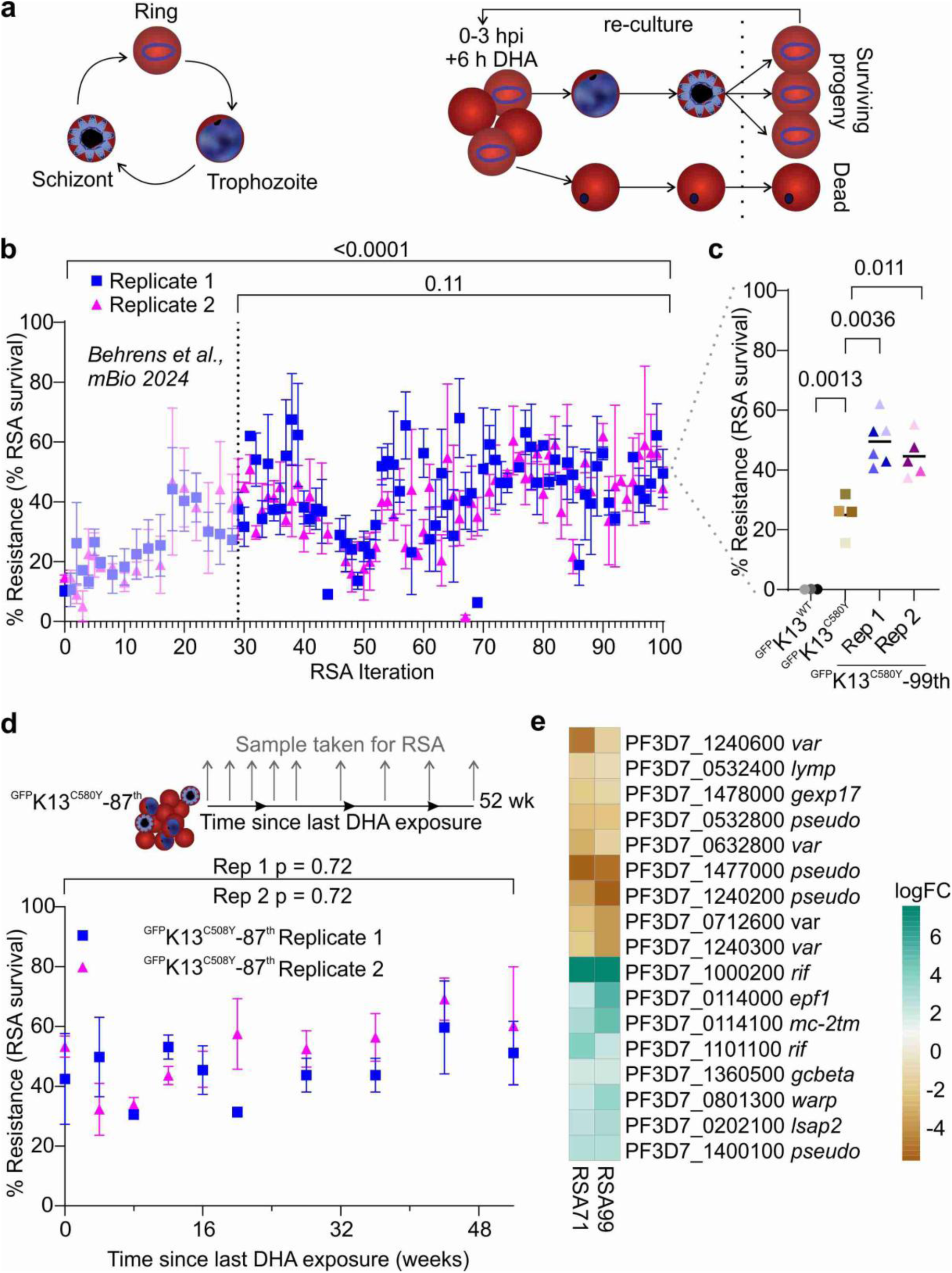
Repeated exposure to DHA selects for stable, hyper ART resistance. **a.** *P. falciparum* blood stage lifecycle (left) and iterative RSA procedure (right). **b.** Increase in resistance (% RSA survival) of ^GFP^K13^C580Y^ with iterative DHA exposure. % survival of 6 h DHA treated parasites compared to control, determined 66 h post treatment for the indicated iteration. Surviving parasites after DHA treatment were recovered and iterative RSAs performed until 100 RSAs. Initial 29 RSAs described previously^2^. Symbols denote mean of 3 independently kept lines per selection replicate with error bars standard deviation (SD). Blue squares replicate 1 and pink triangles replicate 2. **c.** Resistance (% RSA survival) ^GFP^K13^WT^, ^GFP^K13^C580Y^ and survivors of 99 iterative RSAs (^GFP^K13^C580Y^-99^th^) in RSA 100, showing individual experiments and comparison to sensitive and parental lines. Mean ± SD coloured by replicate from (^GFP^K13 ^WT^ black circle n= 3, ^GFP^K13^C580Y^ brown square n= 4 and ^GFP^K13^C580Y^- 99^th^ triangles replicate 1 blue n= 6, replicate 2 pink n= 5). **d.** Resistance (% RSA survival) of hyper resistant parasites over time after cessation of iterative RSAs at RSA 87 (^GFP^K13^C580Y^-87^th^). % RSA survival every 4 weeks and then every 8 weeks. Performed with the lines described in b (two independent selection replicates consisting of 3 lines each). Schematic above. **e.** Heat map showing log Fold Change (logFC) of genes significantly differentially expressed in hyper resistant rings (8-12 hpi) compared to matched ^GFP^K13^C580Y^. ^GFP^K13^C580Y^n = 4, ^GFP^K13^C580Y^-71^st^, n= 3 and ^GFP^K13^C580Y^-99^th^ n= 6. Mean of experiments compared by two-tailed Welch’s t-test.

Mutations in other proteins which ultimately impact haemoglobin digestion can also convey ART-R^2,11–14^, however, the most common mutations are in *k13*, likely as it is functionally required only in ring stages which reduces the fitness cost compared to other endocytosis proteins^2,3,15^. Cell damage repair pathways are also altered in resistant parasites, which in response to ART and, along with parasite genetic background, can contribute to modulating resistance and increased parasite recovery from ART^4,8,12,16–21^. Epigenetic^22^ and epitranscriptomic^23^ changes also contribute to ART-R. However, even in parasite lines harbouring resistance mutations only a portion of parasites survive ART exposure^2,15,24^. The reason for this and what distinguishes survivors from non-survivors, is unclear.

Stochastics in gene expression are a natural cause of phenotypic heterogeneity in diverse organisms^25–28^, with a proposed role in generating population heterogeneity for malaria parasite RBC invasion^29^ and immune evasion^30^ genes. In cancer^31,32^ and bacteria^33–37^ stochastic variation can contribute to drug resistance by generating a diversity of cell states within a genetically isogenic population. These mechanisms usually involve small proportions of dormant or quiescent cells withstanding drug exposure but there is also limited evidence in bacteria that stochastic expression of resistance determinants can influence cell survival without persistence^36,37^. Conversely, the role stochastic cell-to-cell variation plays in drug resistance in malaria parasites is unknown.

As it is only the ring stage that survives ART even in resistant malaria parasites, this resistance is measured through the ring stage survival assay (RSA) where young rings are exposed to a short drug pulse^24^. Exploiting the comparative power of isogenic sensitive (WT), moderately resistant (mutated K13) and “hyper resistant” ART-R parasite lines generated through selection by 99 consecutive RSAs we highlight the central role of K13 as a fulcrum of malaria parasite ART-R and fitness cost. Detailed characterisation revealed that in malaria parasites stochastics of the cellular levels of a single protein, K13, influence cell cycle length and resistance, determining whether an individual cell survives ART or not. This work offers an explanation why and which cells survive ART exposure and shows that the stochastic range of K13 levels in ART-R parasites has shifted to increase the proportion of the population in the ART survivable zone.

## Results

### Repeat DHA pulses drive hyper resistance

Previously, we demonstrated that subjecting ART-R parasites (^GFP^K13^C580Y^) to consecutive RSAs iterations by re-exposure as soon as parasites resurfaced (Fig. 1a), increased parasite resistance after 29 RSAs^2^. Continuation of this selection regime (Fig. 1a, b) for 70 additional RSA cycles (> 18 months of selection) for two independent biological replicates (each with 3 technical replicates constantly kept apart) resulted in resistance that remained around 50 % with minor fluctuations (Fig. 1b). The mean survival of the progeny of parasites which had survived the 99^th^ RSA was 47.3 % in RSA 100 (^GFP^K13^C580Y^-99^th^ parasites, henceforth “hyper resistant parasites”), a significant increase over the parental ^GFP^K13^C580Y^ maintained in long term culture which showed 24.9% survival (Fig. 1c) (“moderately resistant parasites”).

To test the stability of the phenotype we removed parasites from the selection scheme after the 87^th^ RSA iteration when resistance had already plateaued and cultured them without drug pressure. The parasites derived from the 87^th^ RSA maintained the heightened resistance level when cultured in the absence of DHA for one year, indicating the hyper resistance phenotype was stable in *in vitro* culture (Fig. 1d). The generation of stably hyper resistant parasites (^GFP^K13^C580Y^-99^th^) in the same genetic background as sensitive (^GFP^K13^WT^) and moderately resistant (^GFP^K13^C580Y^) parasites provided us with an opportunity to study the properties of the parasites with different ART resistance levels.

### Reduced K13 explains hyper resistance

To characterise the hyper resistant parasites, we carried out transcriptomics with highly synchronous rings, the stage in which ART-R is effective ^24^. In addition to the ^GFP^K13^C580Y^-99^th^ parasites, we also used ^GFP^K13^C580Y^-71^st^ parasites, which were from a point during the selection regime when resistance already had plateaued and showed similar RSA survival levels (Fig. 1b). Most differences in transcript levels present in both hyper resistant parasite lines compared to the parental ^GFP^K13^C580Y^ parasites were from genes encoding exported proteins (15/17, Fig. 1e, Supplementary Fig. 1, Supplementary Data 1), mostly members of variant gene families that frequently show fluctuations between different parasite samples and are not likely to cause a change in resistance. Only two genes encoded non-exported proteins (Fig. 1e, Supplementary Fig. 1, Supplementary Data 1) and both are dispensable for blood stage growth^38,39^, have functions in transmission or mosquito stages^38–41^ and are very lowly expressed in blood stages^38,39^ (Fig. 1e, Supplementary Fig. 1, Supplementary Data 1). Overall, none of the differentially expressed genes were a likely reason for the differences between moderate and hyper resistant parasites. However, the transcriptomics did suggest an age difference between ^GFP^K13^C580Y^ and ^GFP^K13^C580Y^-99^th^ rings despite tight synchronisation (Supplementary Fig 1a).

Whole genome sequencing identified copy number variants and structural variants in ^GFP^K13^C580Y^-95^th^ parasites compared to Pf3D7 WT, but these variants were either present in parental ^GFP^K13^C580Y^ or impacted regions containing primarily exported virulence genes (Supplementary Data 2). No additional mutations to the ^GFP^K13^C580Y^- 29^th 2^ in genes linked to artemisinin resistance were identified (Supplementary Data 2). Mutations in Pf3D7_1329400 (AMP deaminase) and Pf3D7_1205500 (zinc finger protein, putative) that did not fit the criteria for dismissing a potential role mediating changes in resistance were identified. However, experiments validated that these changes were not causal for the hyper resistance (Supplementary Fig. 2).

The cellular abundance of K13 correlates with ART resistance^2^. As the WGS and transcriptomics did not provide any obvious causal changes, we explored whether K13 abundance might be behind the hyper resistance, an idea for which work with the ^GFP^K13^C580Y^- 29^th^ parasites had already indicated a trend^2^. When evaluated using an assay permitting very fine quantification of cellular levels of K13 relative to ^GFP^K13^WT 2,3^, it was revealed that all 6 ^GFP^K13^C580Y^-99^th^ parasite lines had on average less K13 than the parental ^GFP^K13^C580Y^ (Fig. 2a). Considering the average K13 levels in all the 6 separately generated hyper resistant parasite lines the reduction compared to ^GFP^K13^C580Y^ was 12.39 % (45.8 % vs 58.76 % in ^GFP^K13^C580Y^, p= 0.044, Fig. 2a). Hence, despite lacking any further mutations in *k13*, the hyper resistant parasites have even less K13 than the resistant parental ^GFP^K13^C580Y^ parasites. In line with the stability of resistance (Fig 1d) after lifting drug pressure, K13 abundance also remained low across the 6 ^GFP^K13^C580Y^-87^th^ parasites cultured for a year in the absence of DHA pressure (Fig. 2b, Supplementary Fig. 3a). Given the similar resistance and K13 abundance of lines resulting from both biological selection replicates, a single line (^GFP^K13^C580Y^-99^th^ replicate 1A) was used for detailed characterisation.

**Figure 2.**
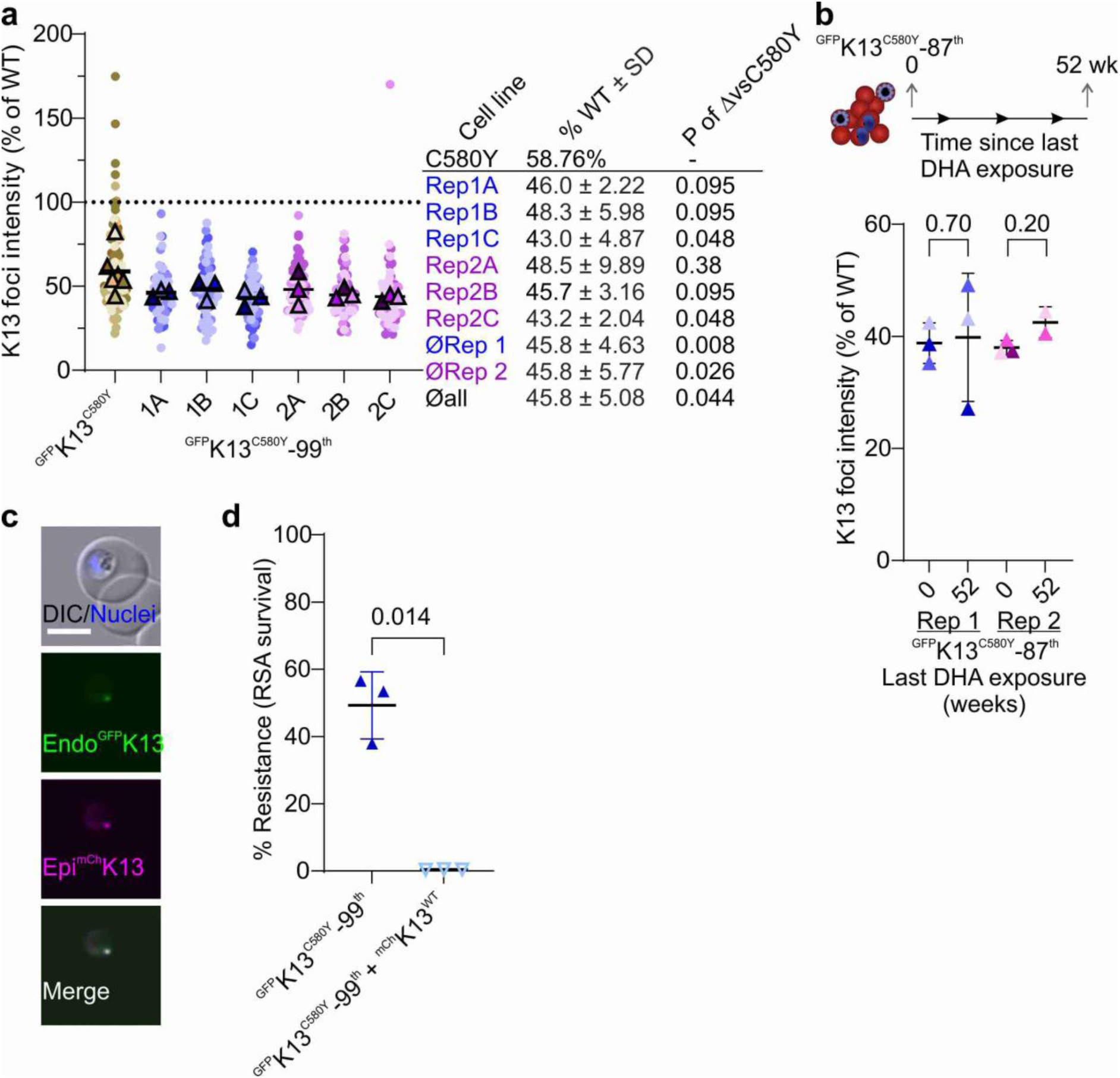
Reduction in K13 abundance explains resistance phenotype of ^GFP^K13^C580Y^- 99^th^. **a.** K13 abundance measured by endogenous GFP fluorescence in ^GFP^K13^C580Y^ and ^GFP^K13^C580Y^-99^th^ rings, normalised to ^GFP^K13^WT^. Individual cells (small symbols) with experimental averages (large symbols) coloured by experiment with mean of average from three independent experiments. Dashed line at 100%. ^GFP^K13^C580Y^ 128 cells, ^GFP^K13^C580Y^-99^th^ 1A 67 cells, 1B 65 cells, 1C 66 cells, 2A 61 cells, 2B 61 cells and 2C 62 cells. Right: Mean K13 experiment average intensity for each line individually and compared means of replicate 1, 2 or all ^GFP^K13^C580Y^-99^th^ lines to ^GFP^K13^C580Y^ by two-tailed Mann-Whitney. **b.** K13 abundance of ^GFP^K13^C580Y^-87^th^ rings after 52 weeks without DHA exposure compared to after 87 RSAs. Abundance measured on one occasion for each selection replicate (average of each of the 3 lines per replicate) from Figure 1b with average of experiment ± SD shown and compared by two-tailed Mann-Whitney test. 20 cells measured for each line per timepoint. Schematic above. **c.** Fluorescence microscopy images of cell line episomally expressing ^mCh^K13^WT^ (epi ^mCh^K13WT) with endogenous ^GFP^K13 in live ^GFP^K13^C580Y^-99^th^ parasites. DIC, differential interference contrast; nuclei (Hoechst), merge (epi ^mCh^K13 and endo^GFP^K13). Representative image, all images (23 cells from 2 independent imaging sessions, Supplementary Fig. 3). Size bar 5 μm. **d.** Resistance (% RSA survival) of ^GFP^K13^C580Y^-99^th^ compared to ^GFP^K13^C580Y^-99^th^ + ^mCh^K13^WT^. % survival compared to control without DHA 66 h after 6 h DHA treatment. Experimental means of three experiments compared by two-sided Welch’s t-test. Mean ± SD and p-value shown.

To test whether the reduction in K13 abundance in ^GFP^K13^C580Y^-99^th^ compared to moderately resistant ^GFP^K13^C580Y^ may be responsible for the increased resistance of these parasites, a second copy of ^mCh^K13^WT^ was episomally expressed in the hyper resistant parasites (Fig. 2c). Additional expression of ^mCh^K13^WT^ reverted ^GFP^K13^C580Y^-99^th^ parasites to complete ART sensitivity (Fig. 2d, reduction from 49.26 % to 0.48 % RSA survival). We conclude that cellular amount of K13 underlies the hyper resistance phenotype and that likely no other major changes that do not impact K13 function or abundance occurred in the serially RSA-selected parasites.

### Hyper resistant parasites have an extended cell cycle

ART-R is known to convey a fitness cost^2,8,42^, with resistant parasites reported to produce fewer progeny^43^, while others reported slowed growth^3,16,44–46^. We therefore assessed growth in the isogenic ART sensitive (^GFP^K13^WT^), moderately resistant (^GFP^K13^C580Y^) and hyper resistant (^GFP^K13^C580Y^-99^th^) parasites lines. First, we evaluated the number of progeny the three parasite lines produced but found no difference (Fig. 3a). Instead, initial experiments using Giemsa smears with synchronous parasites indicated a developmental delay of the resistant parasites (Fig. 3b, two selected time points: mid cycle and end of cycle, both showing that ^GFP^K13^WT^ parasites were fastest, ^GFP^K13^C580Y^ parasites intermediate and ^GFP^K13^C580Y^-99^th^ slowest). To obtain a more quantitative idea about the growth delay, we used flow cytometry and focussed on the time window covering the end of one cycle and the start of the next cycle, measuring changes in the proportions of stages arising (Fig. 3c; Supplementary Fig. 4). These experiments indicated an average cycle length delay of 4 h for the^GFP^K13^C580Y^ parasites (Fig. 3d) and 9.33 h for ^GFP^K13^C580Y^-99^th^ parasites (Fig. 3d) compared to the sensitive ^GFP^K13 ^WT^ parasites. This observation of increased cycle length is consistent with the delayed growth reported for parasites rendered ART-R through K13 mutations^16,45–47^, and as we have used parasite sharing the same genetic background here, this indicates that the growth delay increases with degree of resistance and not genetic background.

**Figure 3.**
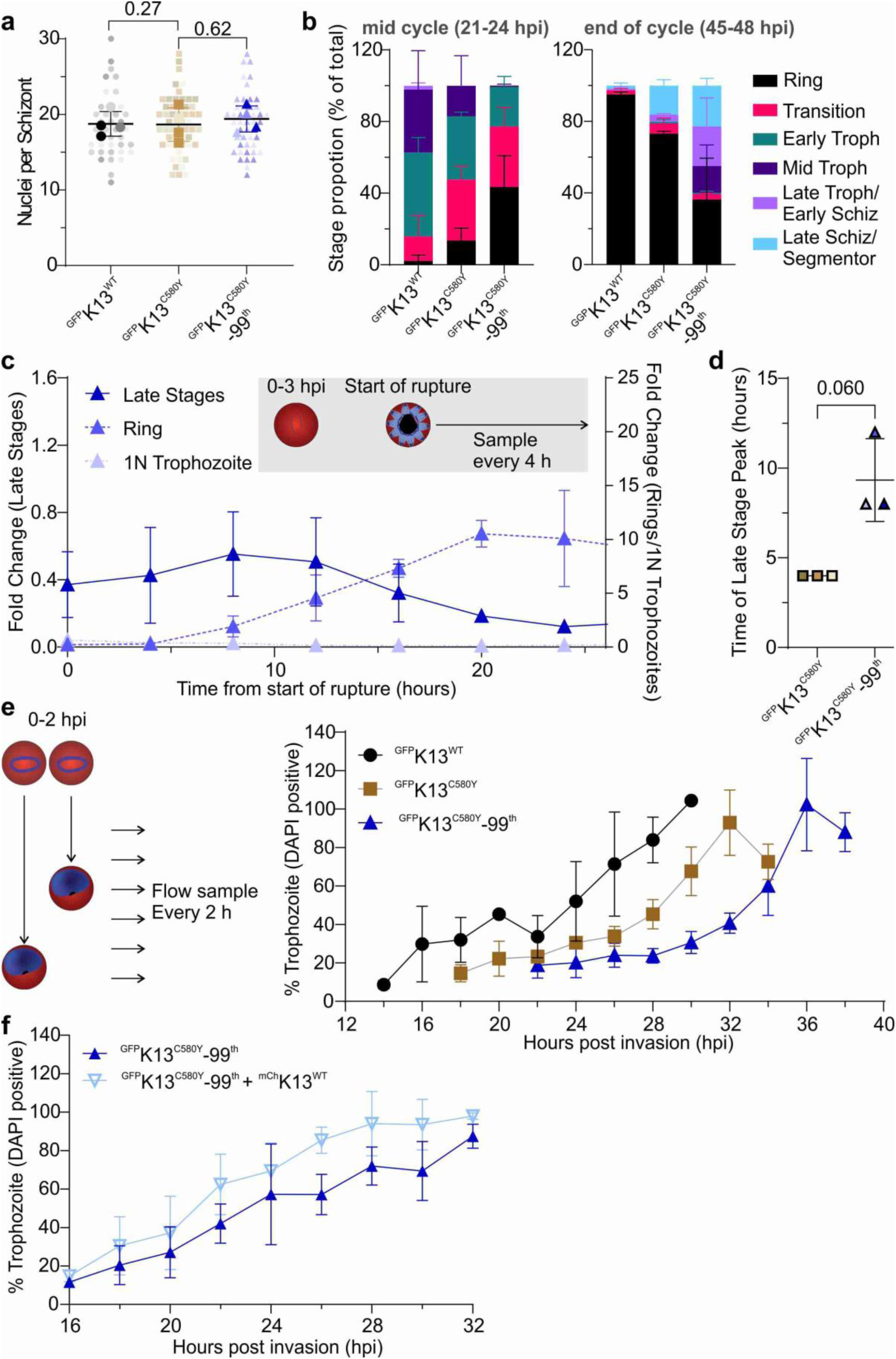
Hyper resistant parasites have altered blood stage cycle progression. **a.** Average schizont nuclei number. Individual schizonts nuclei number (small symbol), experiment average (large symbol; 4 independent experiments differently shaded). ^GFP^K13^WT^ 39 cells, ^GFP^K13^C580Y^ 54 cells and ^GFP^K13^C580Y^-99^th^ 46 cells total. **b.** Stage distribution of 21- 24 hpi (left) and 45- 48 hpi (right) ^GFP^K13^WT^, ^GFP^K13^C580Y^ and ^GFP^K13^C580Y^-99^th^ parasites determined by Giemsa-stained smears in two independent experiments. ^GFP^K13^WT^ 21- 24 hpi = 184 45-48 hpi = 260, 128 and 38 ^GFP^K13 ^C580Y^ 21- 24 hpi = 166 45- 48 hpi = 142 and ^GFP^K13^C580Y^-99^th^ 21- 24 hpi = 141 parasites 45- 48 hpi 104. **c.** Change in late stage, ring and 1 nuclei (1N) trophozoites over time for ^GFP^K13^C580Y^-99^th^. Start of rupture window normalised to ^GFP^K13 ^WT^ assessed in parallel by for 3 independent experiments (Supplementary Fig. 4). Left y axis fold change late stage population, right y axis fold change rings and 1N trophozoites. Grey box: Experiment schematic. **d.** Peak of late stage population for ^GFP^K13^C580Y^ (brown squares) and ^GFP^K13^C580Y^-99^th^ (blue triangles) normalised to ^GFP^K13^WT^. Calculated from b and Supplementary Fig. 4. **e.** Change % trophozoite over time for ^GFP^K13^WT^ (black), ^GFP^K13^C580Y^ (brown) and ^GFP^K13^C580Y^-99^th^ (blue). % of trophozoite (DAPI positive) parasites compared to total (Hoechst and DAPI) determined by flow cytometry over time (left). 3 independent experiments. Left: Experiment schematic. **f.** Proportion of trophozoites overtime for ^GFP^K13^C580Y^-99^th^ compared to ^GFP^K13^C580Y^-99^th^ + ^mCh^K13^WT^ measured by flow cytometry. 4 (^GFP^K13^C580Y^-99^th^) or 5 (^GFP^K13^C580Y^-99^th^ + ^mCh^K13^WT^) independent experiments. Experimental means compared by two-sided Welch’s t-test (a, d). All error bars represent SD.

Given it is ring stages that survive ART and reduction in K13 levels have been shown to extend the ring stage^3,46,47^, we further examined whether a delayed ring stage contributed to the altered cycle length in the ^GFP^K13^C580Y^-99^th^ parasites. Using flow cytometry to follow highly synchronous rings until they had fully transitioned to trophozoites we found that ^GFP^K13^C580Y^- 99^th^ parasites spent ∼ 9 hours longer in ring stage compared to ^GFP^K13 ^WT^ and ∼ 5 hours longer than ^GFP^K13^C580Y^ (Fig. 3e, Supplementary Fig. 4). This confirms the developmental delay suggested by Giemsa smeared parasites (Fig. 3b, mid cycle point). The difference in ring length between ^GFP^K13^C580Y^-99^th^ and ^GFP^K13 ^WT^ parasites (Fig. 3e) matched the total increase in cycle length (Fig. 3c), indicating the delay is attributable to the ring stage alone. Hyper resistant parasites also took longer to develop a second K13 focus likely due to this slowed growth in rings (Supplementary Fig. 4d).

Summarised, these findings show that parasites selected for heightened resistance by serial exposure to DHA have a longer ring phase from which they do not catch up, resulting in an overall increase in cycle length. The ^GFP^K13 ^C580Y^ parasites had an intermediate ring length, again fitting with the difference observed in total cycle length and indicating a proportional relationship of cellular K13 levels, cycle length and resistance.

In addition, these experiments also revealed a reduced stage synchronicity for ^GFP^K13^C580Y^-99^th^ parasites compared to ^GFP^K13^WT^ (Fig. 3c, Supplementary Fig. 4). Despite synchronisation to a 3 h age window in early ring stage, this stage window increased considerably until completion of the cycle with new rings observed around 8 h from start of schizont rupture window (parasite age 44 hpi) while some segmenters remained 20 h after the schizont rupture window had begun (parasite age 56 hpi), compared to ^GFP^K13^WT^ which had negligible late stages by this time (Fig. 3c, Supplementary Fig 4a). Hence, the hyper resistant parasites considerably increased the ring length and show an expanded heterogeneity in cycle length between the different parasites in the population.

Next, we assessed whether the increased cycle length was a result of reduced K13 levels. To test this, we used the ^GFP^K13^C580Y^-99^th^ hyper resistant parasites episomally expressing ^mCh^K13^WT^ (Fig. 2c). These parasites had a reduced length of the ring stage compared to the hyper resistant parasites without extra K13 (Fig. 3f), indicating the elongation of the ring stage seen in our resistant parasites is due to cellular K13 levels. The restriction of the fitness cost to rings is in line with the role of this protein specifically only in ring stages^3^.

Overall, these findings show that in our isogenic parasites cellular K13 levels and cell cycle length (due to the ring stage) correlate with resistance and the rescue by episomal K13 expression shows that K13 levels are the primary causal factor in this.

### Cycle length varies stochastically

While hyper resistant, still only about half of the ^GFP^K13^C580Y^-99^th^ parasites survive the RSA and it is unclear why some cells survive while others die. Given the relationship between ring length and resistance we took advantage of the wide spread in cycle length in the hyper resistant parasites to assess whether the fast-growing parasites differed from the slow-growing parasites. Using highly synchronous ^GFP^K13^C580Y^-99^th^ parasites starting at 0-3 hpi, cells from the faster growing population were isolated at 40-43 hpi, the age at which WT schizonts reach maturity, and a second slower population was collected at 52-55 hpi (Fig. 4a). These isolated populations of fast and slow growers (1^st^ cycle) were allowed to reinvade in a 3 hour window and it was measured how fast these progeny (2^nd^ cycle) reached the schizont stage and ruptured to obtain the cycle length of the 2^nd^ cycle. The progeny from both the fast and slow parasites reached schizont peak at similar times with a similar rate of rupture (Fig. 4b, Supplementary Fig. 5), indicating that both populations had comparable cycle length. There was also no meaningful difference in cycle length between the two populations compared to the average cycle length of ^GFP^K13^C580Y^-99^th^ from Fig. 3 (experiments run in parallel; Fig. 4b, c) with the progeny of the fast population slightly slower than that of the slow population (Fig. 4c; fast 2 ± 2.83 h longer, slow -2 ± 2.83 h than ^GFP^K13^C580Y^-99^th^). The fold increase of ring stages over time was also comparable (Fig. 4d, Supplementary Fig. 5). These findings show that while there is a large variation between individual ^GFP^K13^C580Y^-99^th^ parasites in the time to complete the cycle, fast and slow growers give rise to progeny with the full spectrum of cycle length intrinsic to this cell line, indicating a reset each cycle.

**Figure 4.**
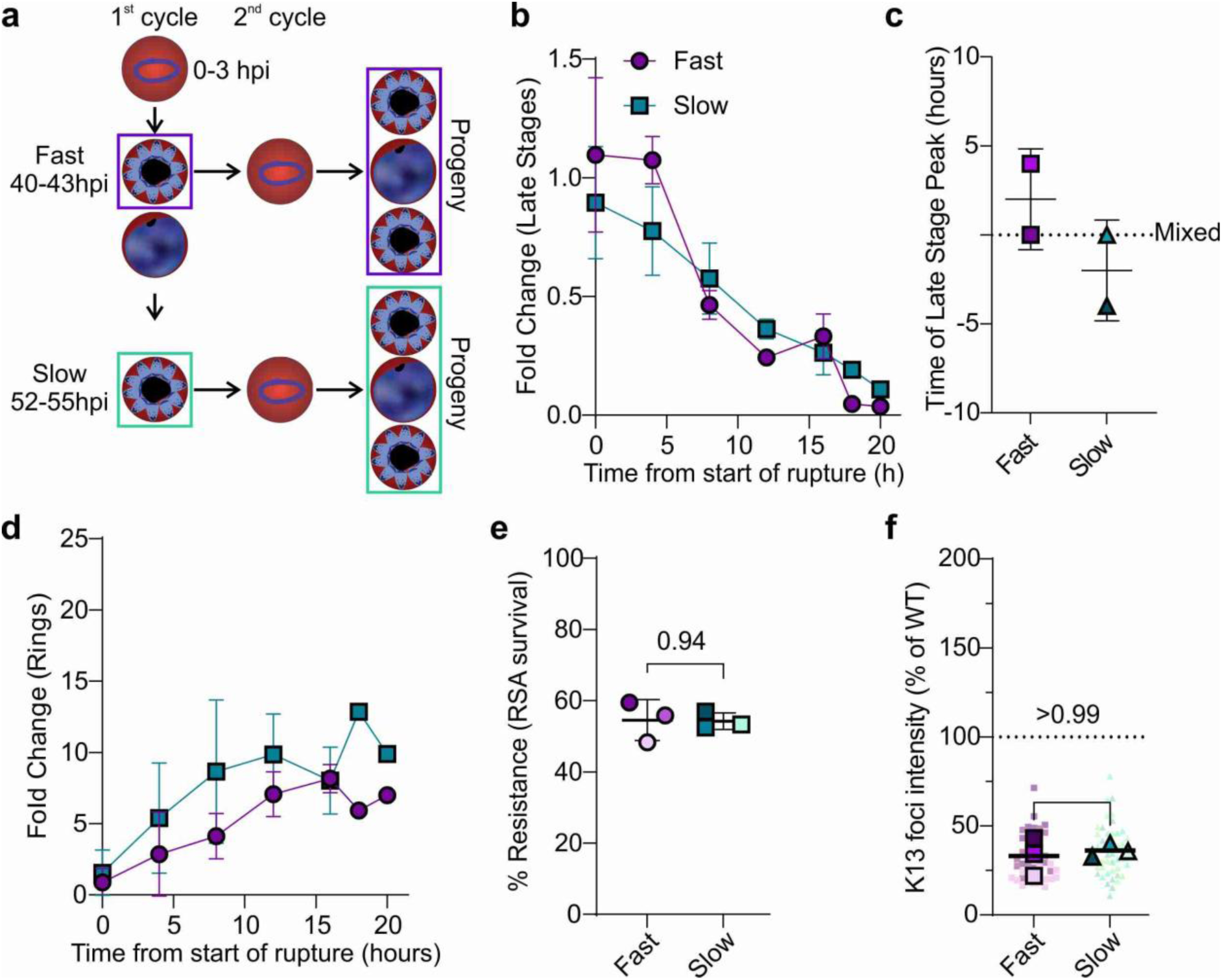
Progeny of fast and slow growing ^GFP^K13^C580Y^-99^th^ display full spectrum of cycle lengths and do not have different ART-R or K13 abundance. **a.** Schematic showing isolation of progeny of fast (40-43 hpi) or slow (52-53 hpi) ^GFP^K13^C580Y-^99^th^. **b.** Change in proportion of late stage progeny (2^nd^ cycle) of fast (purple) and slow (deep teal) growing ^GFP^K13^C580Y^-99^th^ parasites over time determined by flow cytometry. Start of rupture period determined based on the peak of the schizont population of ^GFP^K13^C580Y^-99^th^ in Figure 3 (performed in parallel). **c.** Time difference late stage peak for progeny of the fast (purple) and slow (deep teal) populations was compared, normalised to the peak late stage time point (dashed line “mixed”) of ^GFP^K13^C580Y^-99^th^ not stratified by schizont age (see Methods; Figure 2, performed in parallel). **d.** The fold change in ring (3^rd^ cycle) stage parasites measured concurrently to schizont rupture (b). **e.** Resistance (% RSA survival) of ^GFP^K13^C580Y^-99^th^ parasites resultant from fast (purple) and slow (deep teal) schizonts. % survival compared to control without DHA 66 h after 6 h DHA treatment in RSA. Significance determined by two-sided Welch’s t-test. **f.** Cellular K13 abundance in rings determined by fluorescence intensity of endogenous GFP in progeny from fast (purple) or slow (deep teal) ^GFP^K13^C580Y^-99^th^ normalised to ^GFP^K13^WT^ measured concurrently. Small symbols individual cells (46 cells Fast, 51 cells Slow), large symbols experiment average. Significance determined by two-sided Mann-Whitney test. Error bars represent SD calculated from two experiments (b-d) or three (e, f). Symbols coloured by and representing individual experiments (c, e, f)

We also tested whether fast and slow growers differed in ART resistance. Progeny from fast and slow populations had similar ART-R (Fig. 4e, 54.55 % vs 54.24 % RSA survival), comparable to ^GFP^K13^C580Y^-99^th^ (Fig. 1b). Accordingly, the progeny of slow and fast parasites had comparable K13 levels (Fig. 4f). This demonstrated that ^GFP^K13^C580Y^-99^th^ parasites have increased stochastic variation in cell cycle length that is not individually inherited, resetting each cycle with progeny showing the full spectrum of cycle length, K13 abundance and ART-R.

### Stochastics determine properties of cells

The stochastic nature of gene expression can lead to range of protein levels in individual cells in a genetically clonal population^25–27^. Given the strong relationship between K13 levels and cycle length in our isogenic lines, we can assume that parasites with lower K13 levels would require longer to complete the cycle. To test this assumption on the individual cell level, we measured the cellular abundance of K13 in the hyper ART-R rings and individually tracked these cells by time lapse microscopy until rupture (Fig. 5). Parasites which underwent schizont rupture after 52 hours (corresponding to the average cycle time of WT parasites under time lapse conditions^48^) had higher cellular K13 levels than cells which ruptured later (Fig. 5a). This finding confirms on the single cell level that lower abundance of K13 correlates with an extended cell cycle and supports the variation in cycle length is due to stochastic variation in cellular K13 levels in rings.

**Figure 5.**
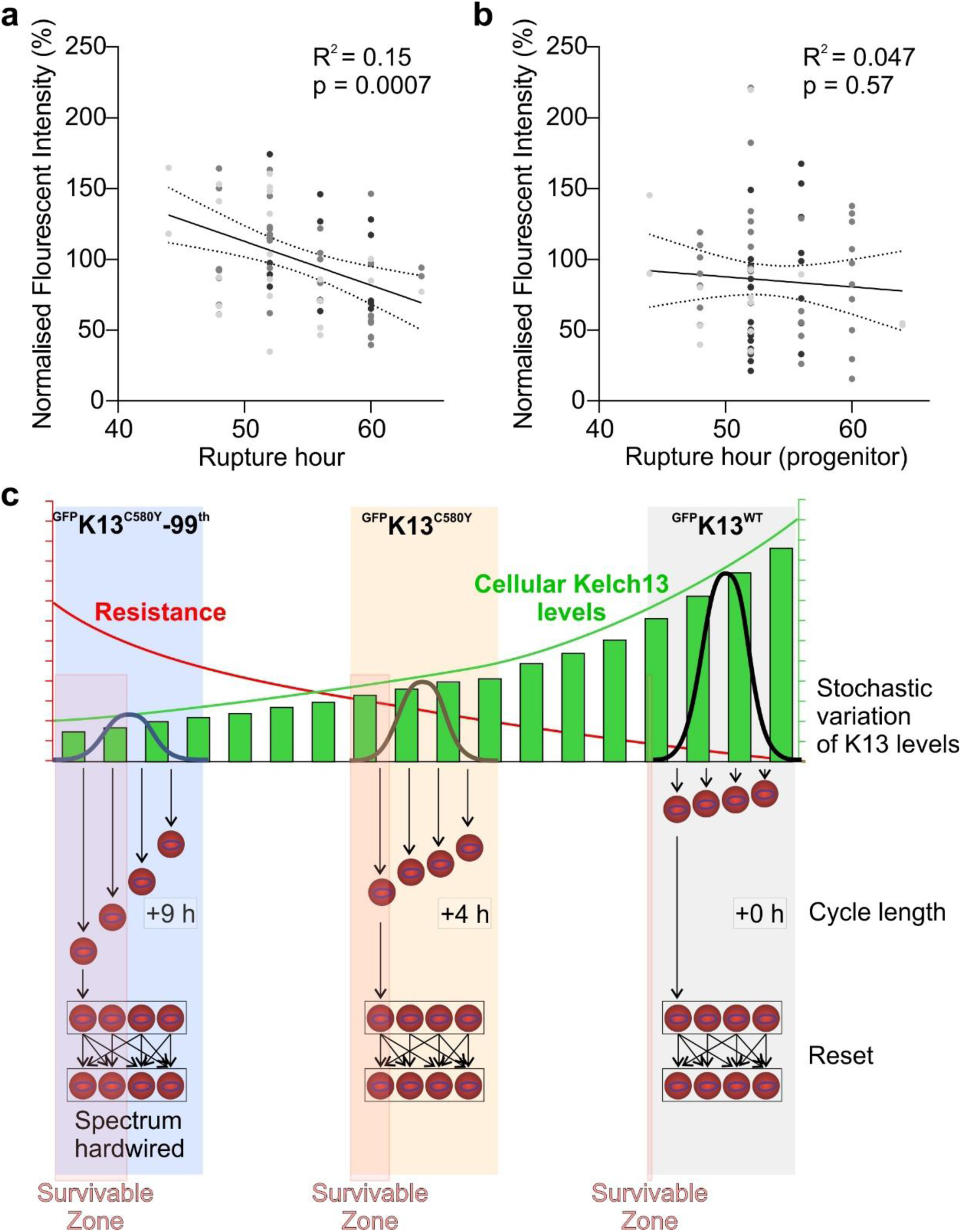
Long term time lapse confirms stochastics determines individual cell fate of hyper resistant parasites. **a.** Individual ^GFP^K13^C580Y^-99^th^ cells were followed by long term time-lapse imaging. K13 abundance of each cell determined in rings and normalised to average experiment fluorescence intensity. Time from experiment start to last time point where a given schizont had not yet ruptured was considered that parasites’ cycle length (Rupture Hour). Each dot represents an individual cell (74 individual cells from 3 independent experiments, marked by different shades). **b.** K13 abundance in progeny compared to rupture hour of progenitor schizont. K13 abundance was normalised to average experiment fluorescence intensity of starting rings. Each dot represents an individual cell (71 individual cells from 3 independent experiments, marked by different shades). R^2^ and significance for Pearson’s correlation between normalised K13 fluorescence intensity and rupture hour. **c.** Schematic highlighting stochastics in K13 abundance (bell curve distribution of K13 levels) and how the resulting cell-to-cell variation results in an increased variation in cycle length and a greater pool of parasites with sufficiently reduced K13 to survive a DHA pulse (survivable zone) specific for each of the cell line (hardwired spectrum) while within population variation is not inherited (reset).

If K13 levels reset each cycle, as indicated by the bulk population experiments (Fig. 4), the progeny arising from the fast and slow cells should again have similar amounts of K13. Assessing K13 levels in our time lapse progeny showed that this was indeed the case (Fig. 5b), confirming resetting of K13 levels in the new generation.

These results indicate that stochastics of K13 levels determine the fate of individual parasites in the population. Parasites with lower K13 levels have a higher chance of surviving the ART pulse at the cost of a slower cycle (Fig. 5c). These survivors give rise to parasites with the full spectrum of K13 levels intrinsic to the specific cell line (Fig. 5c). In resistant parasite this stochastic range has shifted to overall lower K13 levels compared to sensitive parasites, increasing the number of parasites in the range permitting survival of the ART pulse (survivable zone, Fig. 5c).

## Discussion

Here we show that stochastic variation in K13 levels in malaria parasites is directly relevant for parasite fitness and drug resistance. While work examining the average response of bulk populations has established a causal link with RSA survival^2,3,9^, the partial survival phenotype had been puzzling. Based on the causal relationship between K13 abundance, cell cycle length and ART-R in our isogenic lines, we provide evidence that the range of K13 levels determines the average population survival of a given cell line while stochastic expression of K13 determines the survival of individual cells. While the former is inherited and was stable over extended periods of time, the latter is not and each cycle stochastics decide for a cell whether it is above or below the threshold of survival (Fig. 5c).

Previously epigenetics has been explored as an explanation for why only a portion of parasites survive^22,23,49^. Here we have identified a role for gene expression stochastics in ART-R. The influence of cell stochastics in generating heterogeneity within populations is recognised as an important factor in cancer drug resistance^32,50^ and antibacterial resistance^33,35–37^. However, the effect observed here differs from persister population mechanisms seen in bacteria and cancer as it does not found a new population giving rise to resistance. The survivors also do not represent a truly stagnant sub-population waiting out the drug pulse as cellular K13 levels span a continuous range across the parasite population where also the slowest still grow to become trophozoites. Fractional killing of cells has been attributed to multiple factors including non-genetic heterogeneity of cells but typically is complex^51,52^. This work demonstrates that the stochastics of a single protein can impact both fitness and drug resistance, highlighting the central causal role of K13 and providing a comparably simple system showing the phenotypic impact of stochastics.

Our findings have a number of implications. Firstly, while the fitness of individual ART surviving cells will be lower than those parasites with K13 levels above the survivable ART threshold, the progeny of the survivors will again have the full spectrum of the original population before the ART pulse (Fig. 5c). This maximises survival and fitness. Secondly, population measurements, e.g. from ART-surviving parasites, do not necessarily reflect the properties of the cells that survived the ART pulse if not analysed in the same generation. This is relevant for both laboratory and *ex vivo* studies. Thirdly, it can be expected that single-cell analyses of ART-survivors and non-survivors would not find any hard-wired differences. Fourth, the extended ring phase in resistant parasites might indirectly exacerbate resistance by increasing the time of low susceptibility. We would like to stress that this would not be a dormancy related effect, as it simply extends a natural state of the life cycle (extended proportionally to the lowered K13 levels). It is also important to note that this effect would be missed in an RSA, as this assay restricts drug exposure to 6 h. Hence, it does not influence the population RSA survival and hence is not a main driver of ART-R as so far defined, but might be an additional contributor in patients missed by standard *ex vivo* RSAs. Finally, generational resetting of K13 levels, within the population range, may allow parasites to hedge their bets and balance fitness cost and resistance.

In addition to resistance, K13 abundance has also been correlated with fitness cost^2^ and many groups report increased cycle length in K13 mutant parasites^16,46,47^, implying a relationship between K13 abundance and cycle length. Here we explicitly demonstrate that as K13 abundance decreases, average cycle length increases and that this is due to an elongated ring stage. Previous work^45,46^ described resistant parasites with elongated ring stage that “catch-up” to have comparable overall cycle lengths^45,46^. Genetic background is known to impact the resistance^2,8,45^ and fitness cost^2,8^ which might account for this discrepancy, but these studies also used 3D7 parasites^45,46^. Given there was no change in the number of parasite progeny, an increased total cycle length would be more congruent with the known fitness cost of K13 mutations^2,44,47^. One explanation why this was detected here could be that changes in the total cycle length difference may be more obvious in our hyper resistant parasites which have a profoundly elongated cell cycle. Slowed growth of *P. falciparum* harbouring K13 mutations is likely due to decreased amino acid availability, because of reduced amino acid liberation through haemoglobin digestion^3,53^. This is supported by the hyper sensitivity of ART-R parasites to amino acid restriction and increasing amino acid influx into parasites to compensate for overall growth defects in parasites with K13 C580Y^54^.

While we observed a stable reduction in K13 levels in hyper resistant parasites, the cause of this reduction remains unclear. It is possible that changes in the many factors influencing stochastic expression^28^ might also have led to average reduction of K13 levels and the broader cycle length could indicate a loosening of the stochastic rigor indicative of such change. However, given the only ∼10% reduction of K13, and the likely multifactorial influences on stochastics, the magnitude of change in the underlying genes is probably small^28^ and unlikely to be detected in transcriptomics. Alternatively, any change leading to a small reduction in endocytosis (that can be compensated by overexpression of K13), could be a cause of the increased resistance but again these may be too small to be detected by our omics analyses.

The level of stochastic variation in other organisms can vary with pathway^28^. If K13 level stochastics are a bet-hedging strategy, it likely arose from its connection with nutrient uptake not ART-R^49^. Interestingly, codon bias in K13 was proposed to enable bet-hedging through translation regulation^23^, indicating this as one means to obtain differing K13 levels between genetically identical parasites and highlighting K13 expression as a target of non-genetic regulatory processess^23,28,49^.

Both microscopy and flow cytometry-based stage quantifications highlighted a broadening of stage diversity by the end of the blood stage cycle in ^GFP^K13^C580Y^-99^th^ parasites (Fig. 3, Supplementary Fig. 4), suggesting an increase in cycle length variation. Our findings also support an increase in cycle length variation in ^GFP^K13^C580Y^ parasites (Supplementary 4), albeit to a lesser degree. Combined with the observed impact of K13 abundance on cycle length at an individual cell level (Fig 5), this suggest that in populations with minimal K13 protein levels, small cell to cell differences in protein abundance coming from stochastic variation, have a profound impact on cycle lengths and survival under ART. In K13 WT parasites this stochastic variation is less impactful as K13 is replete. Supporting this, parasites harbouring mutations which reduce K13 abundance by approximately 10 % from average WT levels do not cause resistance^2^. Comparatively, a reduction of similar magnitude in parasites with already reduced K13 (13% ^GFP^K13^C580Y^ to ^GFP^K13^C580Y^-RSA99^th^, Fig. 2a) lead to a substantial increase in ART-R which reinforces the evidence for increasing importance of K13 abundance the further it is below WT levels^2^.

Our work demonstrates that 99 cycles of RSAs, an assay where the drug pulse simulates physiological ART action in the host, can increase parasite resistance but only with a corresponding increase in fitness cost. This supports the idea^2^ that the degree to which malaria parasites can become resistant to artemisinin is constrained by fitness cost. Through comparison of isogenic *P. falciparum* parasites with varied ART susceptibility (sensitive, moderately resistant and hyper resistant) we have uncovered a role of stochastic variation in K13 levels in influencing not only individual parasite resistance but also fitness cost. A role for stochastic variation in increasing the proportion of the population of ART-R parasites that are within the “survivable zone” offers an explanation for why some parasites die while others survive under ART in populations uniformly carrying resistance mutations. Since emergence of experimental data showing stochastic cell to cell variation, the impact of these effects has become an appreciated factor in many processes^25–27^. However, in drug resistance the mechanisms usually involve fractional survival^51^ and persister mechanisms^31,32,34,50^. Here we discover that stochastics of a single protein in an eukaryote directly, and likely independent of a persister mechanism, influences phenotypes with implication for drug resistance and parasite fitness.

## Methods and Materials

### Culture of P. falciparum

*P. falciparum* 3D7 parasites were cultured at 5% haematocrit (O+ blood, UKE Hamburg approval 10569a/96-1) in RPMI 1640 (ITW Reagents) with 0.5% Albumax (Life Technologies) at 37 °C under 1 % O_2_, 5 % CO_2_ 94 % N_2_^1^. To generate and maintain transgenic parasite generated through selection linked integration parasites were selected with 4 nM WR99210 (Jacobus Pharmaceuticals) or 2.5 µg/mL BSD (Invitrogen) for episomal carriage and 0.9 µM DSM1 (Sigma Aldrich) or 400 µg/mL G418 (Merck) to obtain integrants as appropriate^2^.

### Generation of gene-edited *P. falciparum* lines

To introduce a second modification into ^GFP^K13^C580Y^ parasites, the pSLI-TGD plasmid^2^ was modified. BSD resistance was amplified using Phusion Polymerase (NEB) and introduced via BamHI/HindIII sites by Gibson. The plasmid was further modified to replace the GFP with a Ty1 tag via BsiWI/SalI. For each desired modification a targeting region was first amplified and introduced into the modified pSLI vector by NotI/MluI Thereafter, the recodonised remainder of the protein of interest (BioCat, Sigma) containing the desired mutation was cloned into the plasmid via MluI/XmaI.

*P. falciparum* ^GFP^K13^C580Y^ parasites were transfected as previously described with episomal expression first selected for by BSD, before integration driven by Neo following the SLI procedure^2^. Integration was confirmed using appropriate primers (Supplementary Data 3) and FirePol polymerase (Solis BioDyne).

### Ring stage Survival Assay (RSA)

To determine the sensitivity of parasites to ART RSAs were performed as previously described^3^. Briefly, young rings (0- 3 h) were exposed to 700 nM DHA (AdipoGen) for 6 h. DHA was then washed off and parasites allowed to grow for a further 66 h after which Giemsa stained (Merck) thin blood smears were made and parasitemia in treated and untreated control determined by microscopy, with smears counted blinded to the identity of the sample. % survival was determined by 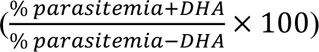 with 1 % the cut-off for resistance^3^.

Selection of hyper resistant parasites through consecutive RSAs was continued as described in Behrens et al.^4^ with two biological selections replicates consisting of 3 technical replicates. DHA exposed cultures from the previous RSA were expanded again until sufficient parasitemia was reached to permit the next RSA. Cryopreserved parasites were revived and selection continued after the 29^th^, 44^th^ and 95^th^ RSA.

### RNA sequencing and differential gene expression analysis

RNA was harvested from 8- 12 h old rings. Ring samples were dissolved in 10 pellet volumes of TRIzol (Life Technologies) then mixed with Chloroform (20% volume) prior to centrifugation at 4 °C for 30 min to separate the top phase containing RNA. RNA was purified using RNeasy Mini Kit (QIAGEN) with additional in-solution DNAse1 (QIAGEN) treatment. Complete removal of gDNA was confirmed by qPCR as previously described^2^ and quality checked by Bioanalyzer 2100 (Aglient) with 6000 Pico Kit. Library preparation was performed with NEBNext Ultra directional RNA library kit (NEB E7760) supplemented with KAPA polymerase and sequenced on DBNSeq platform with 100 bp paired end by BGI Genomics (Hong Kong).

Sequencing data was aligned to the Plasmodium falciparum v61 reference genome^5^ modified to include the recodonised K13 sequence using STAR 2.7.3 aligner^6^ and raw gene counts generated with featureCounts (Rsubread 2.8.2)^7^. To account for staging impact a mixture model was used to estimate stage distribution and perform correction based on ring proportion only prior to differentially expression analysis using Limma/Voom 3.50.3 as has been previously described^8,9^. Samples (1 ^GFP^K13^C580Y^, 3 ^GFP^K13^C580Y^-71^st^) were excluded as proportion of RNA was < 50% ring based on staging. Transcripts per million (TPM) was calculated with Kallisto 0.46.1^10^. Differentially expressed genes were visualised in R using ggplot 2 4.0.2^11^ and pheatmap 1.0.13^12^.

### Whole Genome Sequencing

gDNA was extracted using NEBMonarch Genomic DNA purification kit from 5-10% parasitemia culture. Sequencing using the DNBSEQ PE100 platform and bioinformatic analysis was performed by BGI Tech Solutions (Hong Kong) as previously described^4^. Identified CNV, SV, InDel and SNPs in coding regions in all three technical replicate samples from ^GFP^K13^C580Y^-99^th^ were compared to the previously sequenced parental ^GFP^K13^C580Y 4^. Changes were only considered when found with 100 % frequency in a coding region in all three samples and were absent parental ^GFP^K13^C580Y^ (Supplementary Data 2).

### Determination of length of ring stage by flow cytometry

Due to permeability differences, DAPI does not stain ring stage parasites whereas Hoechst does and this can be used to distinguish rings from trophozoites (manuscript in preparation). Parasites were tightly synchronised by enrichment with 60% Percoll followed by sorbitol 2 hours later^13^. Lines were sampled every two hours. Samples were split and stained with either DAPI alone (trophozoite onwards) or Hoechst and DAPI (all parasites). Stained populations were measured using a UV laser on the LSR ll Flow Cytometer (BD) and paired DAPI only and Hoechst and DAPI samples used to calculate % trophozoites 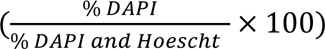. Lines were run in parallel.

### Fluorescence Microscopy

A Zeiss Axio Imager M1 or M2 with a Hamamatsu Orca C4742-95 camera and Zeiss Plan-apochromat 63× oil immersion objectives with 1.4 numerical aperture were used. Long-term time lapse was performed on an Olympus FV3000 with a 1.5 numerical aperture 60X UPlanoApo oil immersion objective. Corel Photo Paint X7 was used to crop images of individual parasites with brightness and intensity adjusted if necessary. Images used for quantification were not adjusted and analysed in ImageJ/Fiji^14, 15^.

### Measurement of K13 fluorescence intensity from bulk populations

Abundance of K13 in rings was determined through measuring the fluorescence intensity of the GFP N-terminally fused on endogenous K13^4,16^. Rings were obtained by synchronisation with 5 % sorbitol 48 h apart. Parasites were returned to culture for a further 2 h before imaging. When assessing abundance in progeny of ^GFP^K13^C580Y^-99^th^ with different cycle lengths parasites were initially synchronised to a 3 h window by Percoll-sorbitol. A second Percoll-sorbitol synchronisation was performed to generate 0- 3 h old rings resultant for ^GFP^K13^C580Y^- 99^th^ schizonts rupturing at 40 hpi (fast) or 52 hpi (slow) ^GFP^K13^C580Y^-99^th^, in parallel with not previously synchronised ^GFP^K13^WT^, and rings imaged immediately. Using the 63x objective cells were selected based on DIC prior to imaging of GFP with the same exposure (200 ms) used for all cell lines. ^GFP^K13^WT^ parasites were always imaged alongside the line of interest and used for normalisation. Foci intensity was measured blinded to sample identity in ImageJ/Fiji^14, 15^ with a minimum of 10 cells per condition per experiment and only cells with a single in focus foci used. Evaluation of distribution for individual cell K13 foci intensity suggested distribution may not be normal and so nonparametric tests were used to compare mean of experiment average.

### Determination of number of K13 foci in rings overtime

Parasites were synchronised to a 3 h window using Percoll- sorbitol and split into 3 identical dishes. At 14- 17 hpi, 18-21 hpi or 22-25 hpi one dish was used for imaging. 15 cells minimum were imaged by fluorescence microscopy for each line at each timepoint and number of K13 foci per cell counted blinded.

### Nuclei number in mature schizonts

Parasites were synchronised to a 4 h window by Percoll-sorbitol and 1 µM Compound 2^17^ was added to 38-41 hpi schizonts and incubated for 6 h at 37 °C. Compound 2 was then washed out and parasites stained with Hoescht (50 ng/uL) for 15 min prior to imaging by fluorescent microscopy. A minimum of 15 cells were captured per line per replicate.

### Measurement of cell cycle stochastics

Two approaches were taken to confirm cell cycle length and dynamics. First, stage distribution was evaluated based on Giemsa-stained thin blood smears of parasites with the 3 h window generated by Percoll- sorbitol with smears made from the same culture at 21 – 24 hpi and 45- 48 hpi.

A second flow cytometry-based approach was also used to study shifts in cell cycle length. ^GFP^K13^WT^, ^GFP^K13^C580Y^ and ^GFP^K13^C580Y^-99^th^ parasites were synchronised to a 3 h window by Percoll- sorbitol with total parasitemia determined at 24 hpi by flow cytometry. Starting at 36 hpi the portion of ring, 1 N trophozoites at >2 N parasites (Late stages) was determined every 4 h by flow cytometry. The fold change in stage populations was calculated against the starting parasitemia for each line with the start of rupture period defined as the timepoint where the ^GFP^K13^WT^ late stage population was maximal. Changes in each population (Ring, 1N Trophozoite, Late stage) over the rupture window was visualised and used to determine the timepoint at which the late stage population was maximal representing the average cycle length. The slope of the late stage population from start of rupture window until < 10% of starting parasitemia was calculated by linear regression.

To determine whether progeny showed similar dynamics the above experiment was modified with tight synchronisation (0-3 h window, 1^st^ cycle) performed prior to tight synchronisation of progeny (2^nd^ cycle) from “fast” growing parasites (schizont isolated at 40 hpi) and “slow” (schizont isolated at 52 hpi) parasites. These resultant progenies (end of 2^nd^ cycle, start of 3^rd^ cycle) were tracked as above, except normalisation was too ^GFP^K13^C580Y^-99^th^ parasites which had not undergone the initial Percoll-sorbitol synchronisation (mixed).

### Long term timelapse of ^GFP^K13^C580Y^-99^th^ parasites and measurement of K13 intensity from individual cells

Tightly synchronised (0-3 h) ^GFP^K13^C580Y^-99^th^ parasites were coated onto µ-Dish 35 mm, high (81151, ibidi) which had been coated with conA (Sigma, C0412) as described previously^13^. Imaging was performed on Olympus FV3000 with heated imaging chamber at 37 °C. To determine average K13 intensity of individual rings 5 z stack (4 z stacks third replicate, 0.32 µm steps between confocal planes) were acquired 30 minutes apart. To track parasite growth subsequent images were taken at 24, 40, 44, 48, 52, 60, 64 and 68 hpi.

The average K13 foci intensity in each ring was determined by measuring the K13 fluorescence intensity in maximum intensity projection after background subtracted. Measurements from the 5 initial z-stacks (4 in third replicate) for each ring were then averaged to get that rings’ average K13 foci intensity. This was then normalised to the mean intensity of all K13 ring measurements from that experiment to control for experiment to experiment variability. Measurements were excluded if inspection of z stack indicated the entire K13 signal was not captured. For assessing K13 intensity in progeny rings only 1 time point was used and normalisation was to the average K13 intensity determined at the start of that experiment. Only rings that could confidently be identified as progeny of the schizont of interest were included. The last time point prior to schizont rupture was taken as the cell cycle length as rupture occurred before the next time point. RBCs with double infections and parasites which had two K13 foci in rings or failed to rupture within the experiment time frame were excluded from analysis. Image analysis was performed in ImageJ/Fiji^14, 15^.

### Statistical Analysis

Statistical evaluation of results was performed in either GraphPad Prism 11.0.2 or R 2022.02.02. Type of statistical test performed, with replicate information, indicated in Figure legend. Normal distribution assumed unless otherwise stated.

## Data and code availability statement

All data supporting the findings of this study are available within the paper and supplementary information.

## Acknowledgements

We thank Jakobs Pharmaceuticals for supplying WR99210. The authors also thank Anna Bachmann for advice and helpful discussion about transcriptomic analysis. We thank the BNITM Flow Cytometry facility for instrument support.

## Author Contributions

IGH, PL-B, SS, LP and VH performed experiments. IGH and PL-B performed formal data analysis and prepared Figures. TS obtained funding. IGH and TS were responsible for experimental design. IGH and TS wrote the manuscript draft. All authors approved the manuscript.

## Additional information

Supplementary Information is available for this paper.

