## Supplemental figure and legends for "Kelch13 stochastics determine drug survival in resistant malaria parasites"

### Supplementary Figure Legends

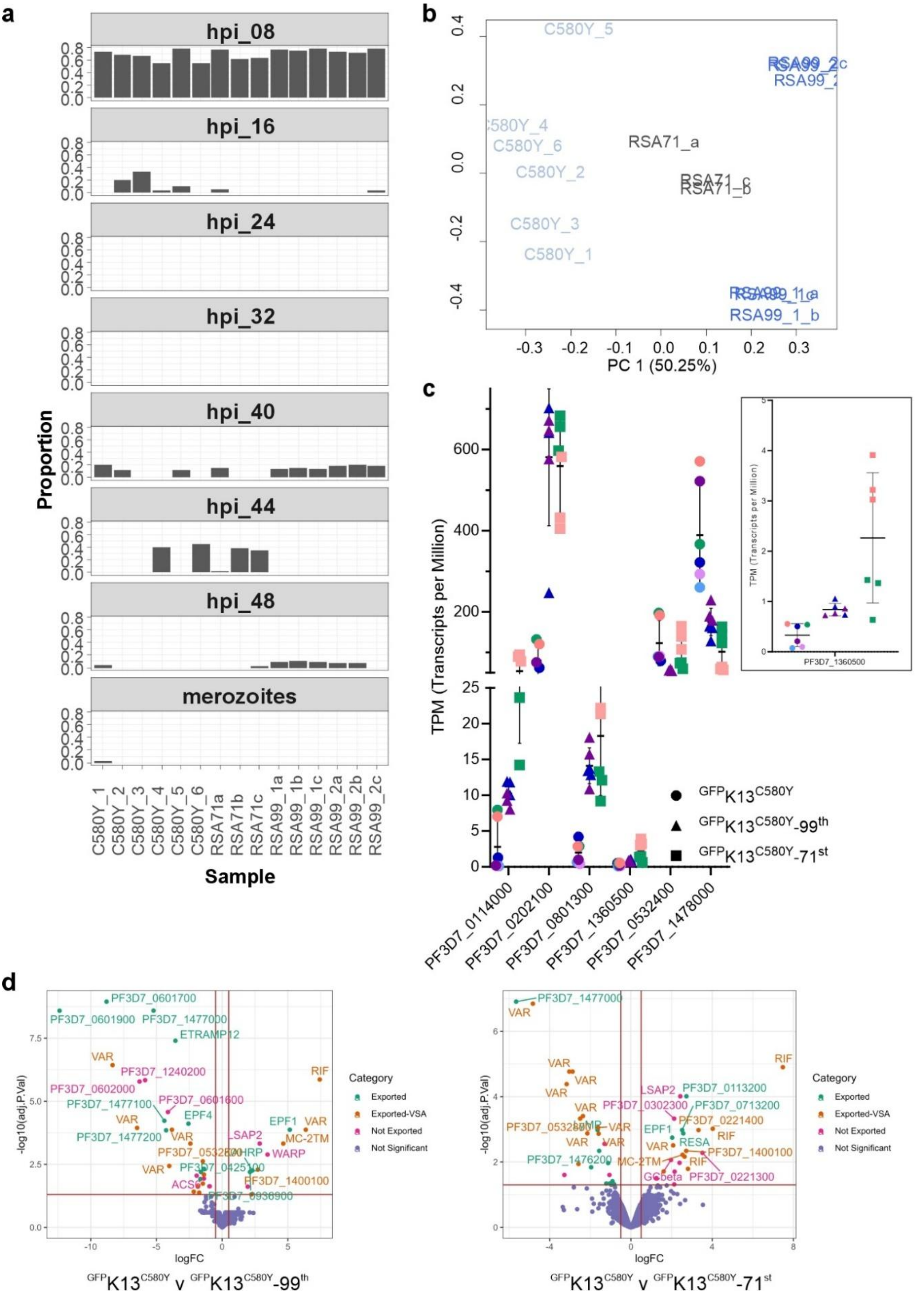


**Supplementary Figure 1. Stage distribution and differentially expressed genes in hyper resistant parasites.**

Transcriptome of 8- 12 h old rings from parental$^{GFP}$K13$^{C580Y}$n = 4 were compared to $^{GFP}$K13$^{C580Y}$-71$^{st}$ n= 3 and $^{GFP}$K13$^{C580Y}$-99$^{th}$ n = 6.

**a**. Proportion of each stage based on transcriptomic profile of sample compared to time course of Pf3D7 over the blood stage (Wichers et al., *mBio* 2019).

**b**. Principle Component Analysis for transcriptomic samples after application of the mixture model [52] to correct for stage differences. Samples from $^{GFP}$K13$^{C580Y}$-71$^{st}$ dark grey, $^{GFP}$K13$^{C580Y}$-99$^{th}$ blue and $^{GFP}$K13$^{C580Y}$ grey.

**c**. Transcripts per million (TPM) of genes differentially expressed between $^{GFP}$K13$^{C580Y}$ (circles) and hyper resistant parasites ($^{GFP}$K13$^{C580Y}$-71$^{st}$ square and $^{GFP}$K13$^{C580Y}$-99$^{th}$ triangle), excluding *Pf*EMP1s, rifins and pseudogenes. Insert top right: Pf3D7_1360500. Samples harvested in parallel are represented in the same colour.

**d**. Differential expression of genes in 8-12 h old rings in $^{GFP}$K13$^{C580Y}$ compared to $^{GFP}$K13$^{C580Y}$-99$^{th}$ (left) and $^{GFP}$K13$^{C580Y}$-71$^{st}$ (right). Significantly differentially expressed genes of exported proteins labelled green, genes of exported proteins which are members of a Variant Surface Antigen (VSA) multigene family labelled orange and genes differentially expressed but not coding exported proteins are labelled pink. Not significant labelled purple. Red lines mark cut off of logFold Change (logFC) < +/- 0.5, p adjusted value < 0.05. Genes labelled when logFC < +/- 1.5, p adjusted value < 0.01 for ease of reading with either abbreviation or where no abbreviation exists Gene ID.

a

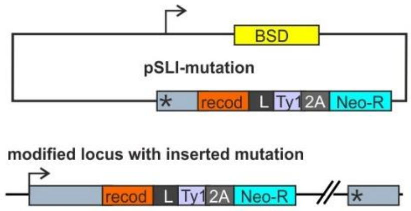


b

| GeneID | Name | Mutation | Comment |
|---|---|---|---|
| PF3D7_0519300 | protoheme IX farnesyl-transferase | c.1096T>C p.Phe366Leu | Mutation not in conserved functional residues (Gurge et. al Scientific Reports 2019). Mutation in parental $^{GFP}K13^{C580Y}$ (p.Asn2Lys) not present in hyper resistant and thus protein excluded from further validation. |
| PF3D7_0910200 | conserved Plasmodium protein, unknown function | c.7501A>T p.Asn2501Tyr | Excluded from further validation as *P. falciparum piggybac* screen (Zhang et al., Science 2018) indicates gene is not essential and disruption does not convey strong fitness cost (MFS score). |
| PF3D7_0929400 | high molecular weight rhoptry protein 2 | c.3773T>G p.Ile1258Ser | Excluded from further validation as mutation in unstructured region and RhopH2's known trophozoite and late stage function with no current indications of ring role. |
| PF3D7_1205500 | zinc finger protein, putative | c.1694G>C p.Arg565Thr | Mutation had no impact on % survival in RSA, this work. |
| PF3D7_1329400 | AMP deaminase | c.2044A>T p.Asn682Tyr | Mutation had no impact on % survival in RSA, this work. |
| PF3D7_1357800 | T-complex protein 1 subunit delta | c.57A>T p.Gln19His | Excluded from further validation as mutation only detected in portion of $^{GFP}K13^{C580Y}$-71$^{st}$ transcripts which have comparable resistance to $^{GFP}K13^{C580Y}$-99$^{th}$ parasites. |

c

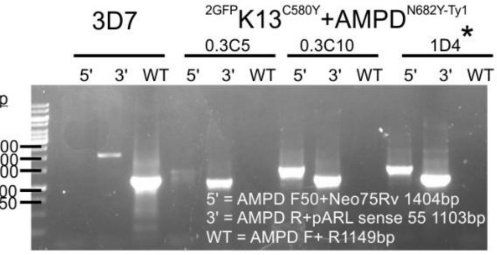


d

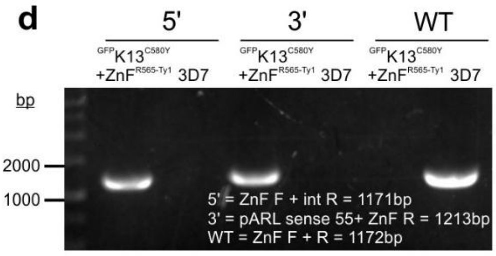


e

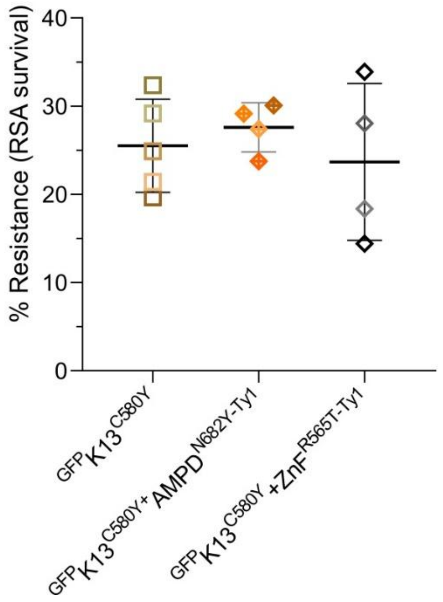


**Supplementary Figure 2. Introduction of selected mutations into $^{GFP}K13^{C580Y}$ to test impact on ART-R.**

**a**. Schematic showing the modification of the second locus in the parental $^{GFP}K13^{C580Y}$. The region starting from just prior to the identified mutation to the end of the protein was replaced with a recodonised segment containing the mutation.

**b**. Table describing location and change of for all SNP that were present in the genome of all 3 hyper resistant samples and not in $^{GFP}K13^{C580Y}$. Full list of SNPs and criteria for validation in Supplementary Data 2.

**c**. Correct integration was confirmed by PCR from gDNA shown in $^{GFP}K13^{C580Y}$+$AMPD^{N682Y\text{-}Ty1}$ or **d.** $^{GFP}K13^{C580Y}$+$ZnF^{R565T\text{-}Ty1}$. For $^{GFP}K13^{C580Y}$+$AMPD^{N682Y\text{-}Ty1}$ 3 lines were generated with 1D4 used for all experiments.

**e**. Identified mutations in PF3D7_1329400 (AMPD) and PF3D7_1205500 (ZnF) were individually introduced into parental $^{GFP}K13^{C580Y}$ and RSAs performed to test impact on ART-R. $^{GFP}K13^{C580Y}$ n= 5, $^{GFP}K13^{C580Y}$+$AMPD^{N682Y\text{-}Ty1}$ n= 4 and $^{GFP}K13^{C580Y}$+$ZnF^{R65T\text{-}Ty1}$ n= 4. Experiments shaded by colour, mean % survival compared by two-sided Welch's t-test.

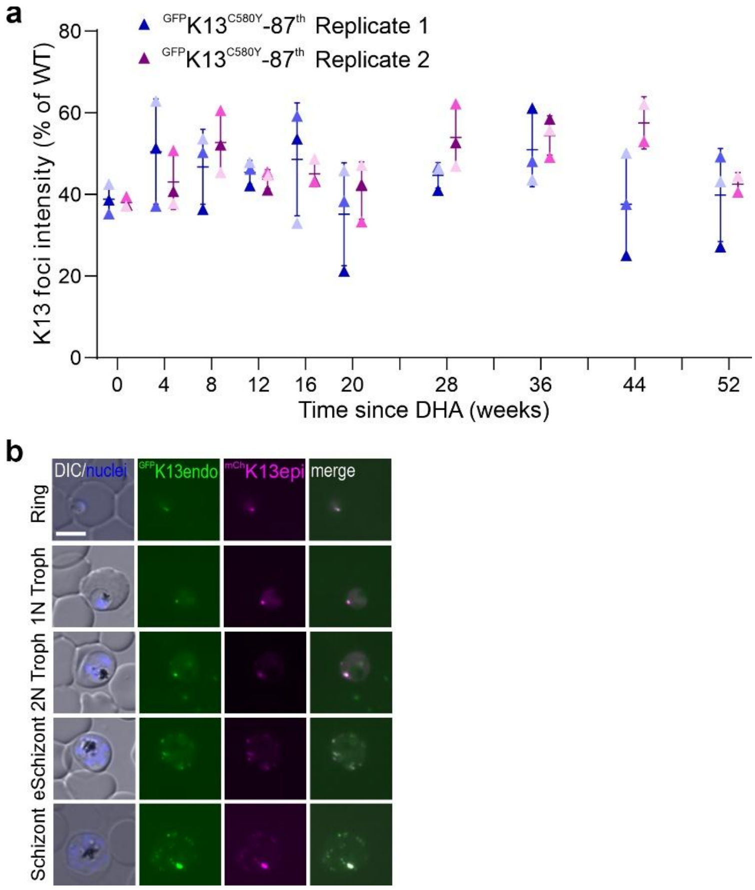


**Supplementary Figure 3. K13 abundance and overexpression in hyper resistant parasites.**

**a**. Endogenous GFP fluorescence intensity of the single K13 foci in ring, normalised to $^{GFP}$K13$^{WT}$, of $^{GFP}$K13$^{C580Y}$-87$^{th}$ after the cessation of iterative RSA pulses. Changes in K13 abundance were initially measured every 4 weeks and then every 8 weeks. Abundance was measured on one occasion for each of the replicates (3 lines per replicate) from Figure 1b. Average K13 foci intensity of 20 cells shown from each line coloured by intensity. Short horizontal line represents across the three selected lines for that time point with error bars SD. **b**. Live microscopy of different stages $^{GFP}$K13$^{C580Y}$-99$^{th}$ + $^{mCh}$K13$^{WT}$ parasites. DIC and Hoescht stained nuclei (blue) represented in merge, endogenous $^{GFP}$K13 in green, episomally expressed $^{mCh}$K13 in magenta with final column containing merge of both K13 signals (representative images from 23 cells from two independent occasions). Size bar 5 µm. Example image displayed in Figure 2.

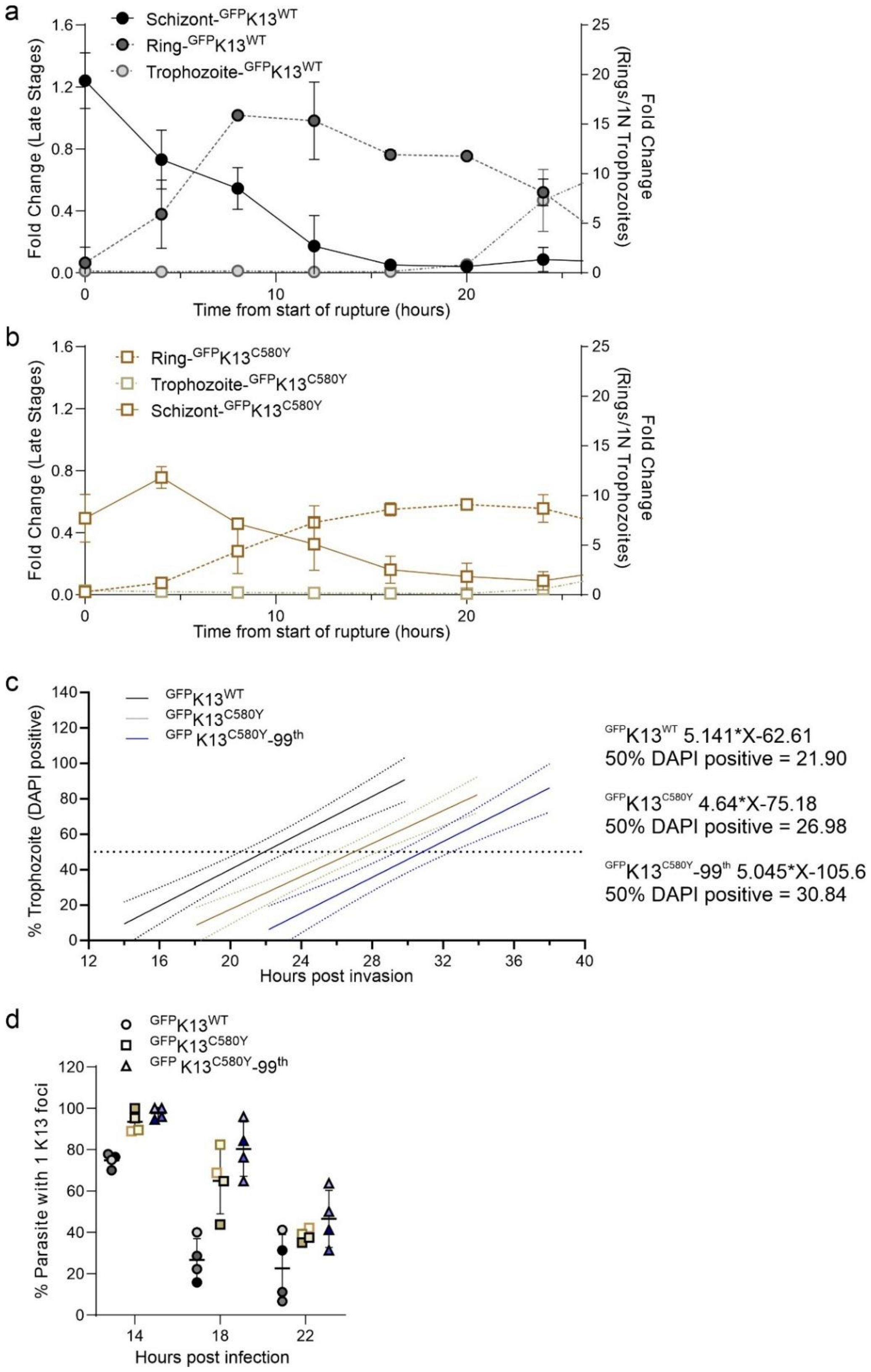


**Supplementary Figure 4. Phenotypic characterisation of hyper resistant parasites.**

**a-b.** The fold change in rings, 1 nuclei trophozoites and late stages (>2 nuclei) over time for$^{GFP}$K13$^{WT}$ **a.** and $^{GFP}$K13$^{C580Y}$ **b.** compared to starting parasitemia. All populations determined by flow cytometry in parallel. Left y axis gives fold change for late stage population while right y axis gives fold change for rings and 1 nuclei trophozoites. Mean from 3 experiments shown with error bars representing SD.

**c**. Linear regression of rate of ring to trophozoite transition for $^{GFP}K13^{WT}$ (black), $^{GFP}K13^{C580Y}$ (brown) and $^{GFP}K13^{C580Y}$-99$^{th}$ (blue) parasites calculated from Figure 2C. Dotted lines represent 95% confidence interval. Slope of linear regression was used to calculate the time at which 50 % of the population reach trophozoite (right).

**d**. The number of K13 foci over time in $^{GFP}K13^{WT}$, $^{GFP}K13^{C580Y}$ and $^{GFP}K13^{C580Y}$-99$^{th}$ determined by live fluorescence microscopy. The percentage of parasites with only once focus compared to the total number of cells for each line at three time points. A minimum of 15 cells was imaged per line per time point and foci counting conducted blinded. Experiments coloured by shade.

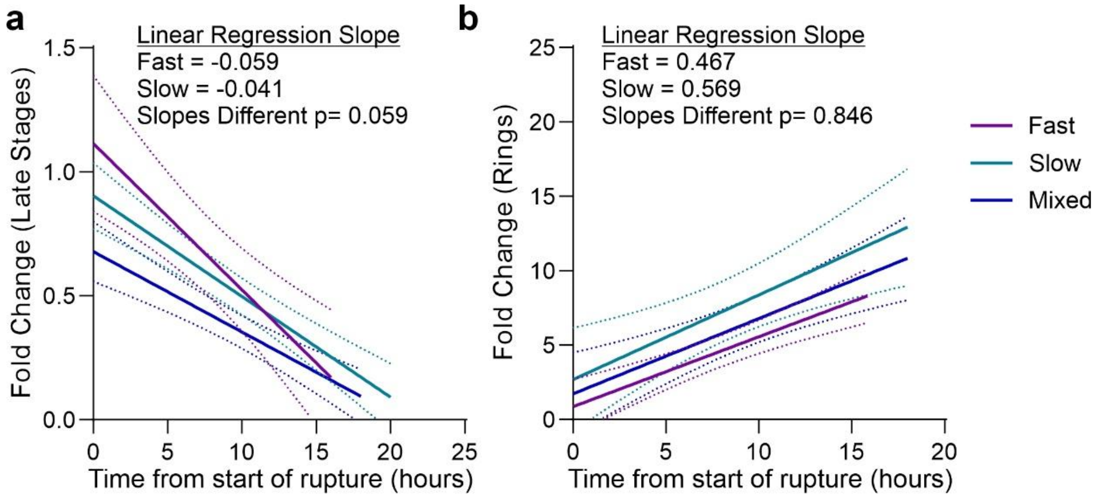


**Supplementary Figure 5. Linear regression of changes in late stage and ring populations over time for progeny from fast and slow growing $^{GFP}$K13$^{C580Y}$-99$^{th}$.**

**a**. Rate of schizont rupture in Figure 4 determined as the slope of the linear regression as a proxy for the spread of ages present within the population. Calculation also shown for mixed ($^{GFP}$K13$^{C580Y}$-99$^{th}$ not stratified by age, Figure 4), which was used to establish when the rupture window started and was performed in parallel.

**b**. Linear regression Figure displaying increase in ring population from start of rupture until time point with highest ring stage parasitemia.

Solid lines represent mean with 95% confidence interval as dashed lines. Slope of each linear regression calculated and compared by two-tailed test. Two independent experiments.

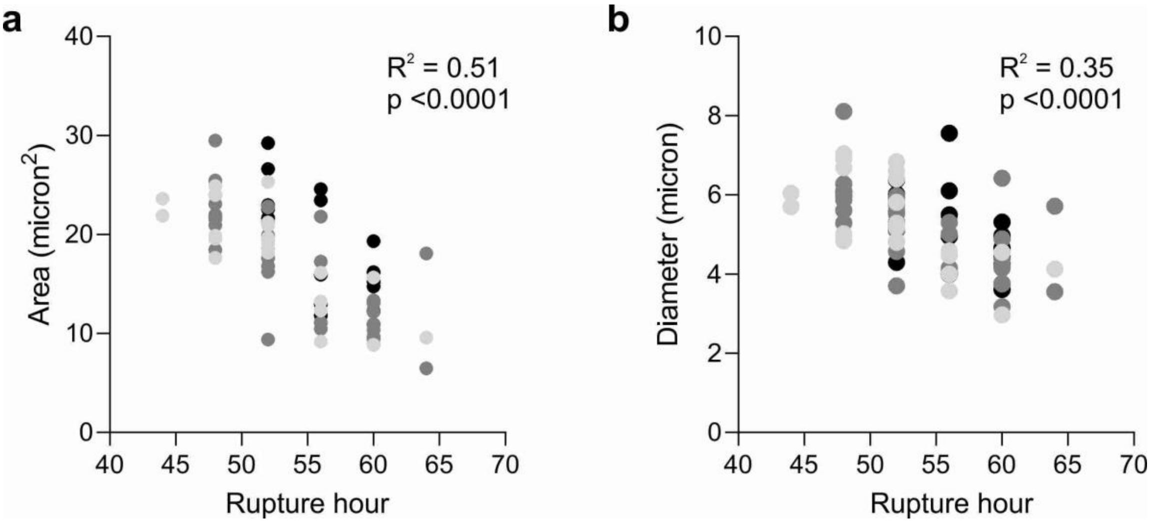


**Supplementary Figure 6. Relationship between cell size and hour of rupture for $^{GFP}$K13$^{C580Y}$-99$^{th}$.** Area **a.** and diameter **b.** of individual parasites at 40 hpi compared to time (hpi) of rupture for hyper resistance $^{GFP}$K13$^{C580Y}$-99$^{th}$ analysed in Figure 5. Each dot represents one cell, with same colour used for each of the three experiments. 74 cells total. Pearson's correlation performed to identify meaningful correlation.
